# G protein-coupled estrogen receptor regulates heart rate by modulating thyroid hormone levels in zebrafish embryos

**DOI:** 10.1101/088955

**Authors:** Shannon N Romano, Hailey E Edwards, Xiangqin Cui, Daniel A Gorelick

## Abstract

Estrogens act by binding to estrogen receptors alpha and beta (ERα, ERβ), ligand-dependent transcription factors that play crucial roles in sex differentiation, tumor growth and cardiovascular physiology. Estrogens also activate the G protein-coupled estrogen receptor (GPER), however the function of GPER *in vivo* is less well understood. Here we find that GPER is required for normal heart rate in zebrafish embryos. Acute exposure to estrogens increased heart rate in wildtype and in ERα and ERβ mutant embryos but not in GPER mutants. GPER mutant embryos exhibited reduced basal heart rate, while heart rate was normal in ERα and ERβ mutants. We detected *gper* transcript in discrete regions of the brain and pituitary but not in the heart, suggesting that GPER acts centrally to regulate heart rate. In the pituitary, we observed *gper* expression in cells that regulate levels of thyroid hormone triiodothyronine (T3), a hormone known to increase heart rate. GPER mutant embryos showed a mean 50% reduction in T3 levels compared to wildtype, while exposure to exogenous T3 rescued the reduced heart rate phenotype in GPER mutants. Our results demonstrate that estradiol plays a previously unappreciated role in the acute modulation of heart rate during zebrafish embryonic development and suggest that GPER regulates basal heart rate by altering total T3 levels.

## INTRODUCTION

Zebrafish are an established model for human cardiovascular development and function (1) with conserved estrogen signaling (2–4). While studying the function of ERα (*esr1*) in zebrafish embryonic heart valves (5, 6), we serendipitously observed that estrogen receptor modulators caused acute changes in heart rate. Estrogens bind two classes of receptors: nuclear hormone receptors (ERα, ERβ) that are ligand-dependent transcription factors (7), and the G protein-coupled estrogen receptor (GPER, also known as GPR30), an integral membrane protein (8, 9). It has been difficult to tease apart to what degree ERα and/or ERβ are involved in regulating GPER function *in vivo*. The observations that ERα can directly activate G proteins in cultured cells (10–13) and that GPER coimmunoprecipitated with ERα in tumor cells (14) has been used to argue that either GPER is dispensable for estrogen-dependent signaling or that GPER mediates interactions between ERα and G proteins (15). Studies using GPER-deficient mice implicate GPER in ventricular hypertrophy (16), regulation of blood pressure and vascular tone (17, 18) and atherosclerosis progression (19), but whether nuclear ER signaling is required for GPER function in these contexts is unknown. Additionally, these studies examined GPER function in adult animals, while the role of GPER during embryonic development is not well understood. Here we use zebrafish embryos, an established model of human development, to reveal a new function for GPER during cardiovascular development.

Estrogen signaling often differs between males and females. However, zebrafish embryos and larvae are bipotential hermaphrodites that have not begun to sexually differentiate before approximately 10 days post fertilization (dpf) (20), meaning that estrogen levels are uniform between age-matched embryos. Additionally, zebrafish embryos develop outside of the mother and not within a confined space, such as the uterus. Therefore, zebrafish embryos are not subject to local estrogen concentration gradients, as has been reported to occur in rodents depending upon their position *in utero* and their proximity to embryos of the same or opposite sex (21, 22). These developmental traits make zebrafish a powerful model to study how sex hormone signaling influences the formation and function of non-gonadal tissues. Using complementary genetic and pharmacologic approaches, we sought to characterize how estradiol regulates heart rate and determine to what extent each estrogen receptor mediates estradiol-dependent changes in heart rate in zebrafish embryos.

## RESULTS

We exposed 49 hour post fertilization (hpf) embryos to 17β-estradiol (estradiol) and assayed heart rate following one hour exposure. We found that estradiol exposure caused an approximately 20% increase in heart rate (Fig. 1, mean difference in heart rate between estradiol and vehicle exposed embryos was 26.51 ± 3.36 (standard error) beats per minute (bpm)). Exposure to progesterone, a structurally similar steroid sex hormone, had no effect on heart rate (Fig. 1, mean difference in heart rate 2.31 ± 6.54), suggesting that the effects on heart rate were specific to estrogens.

**Figure 1.**
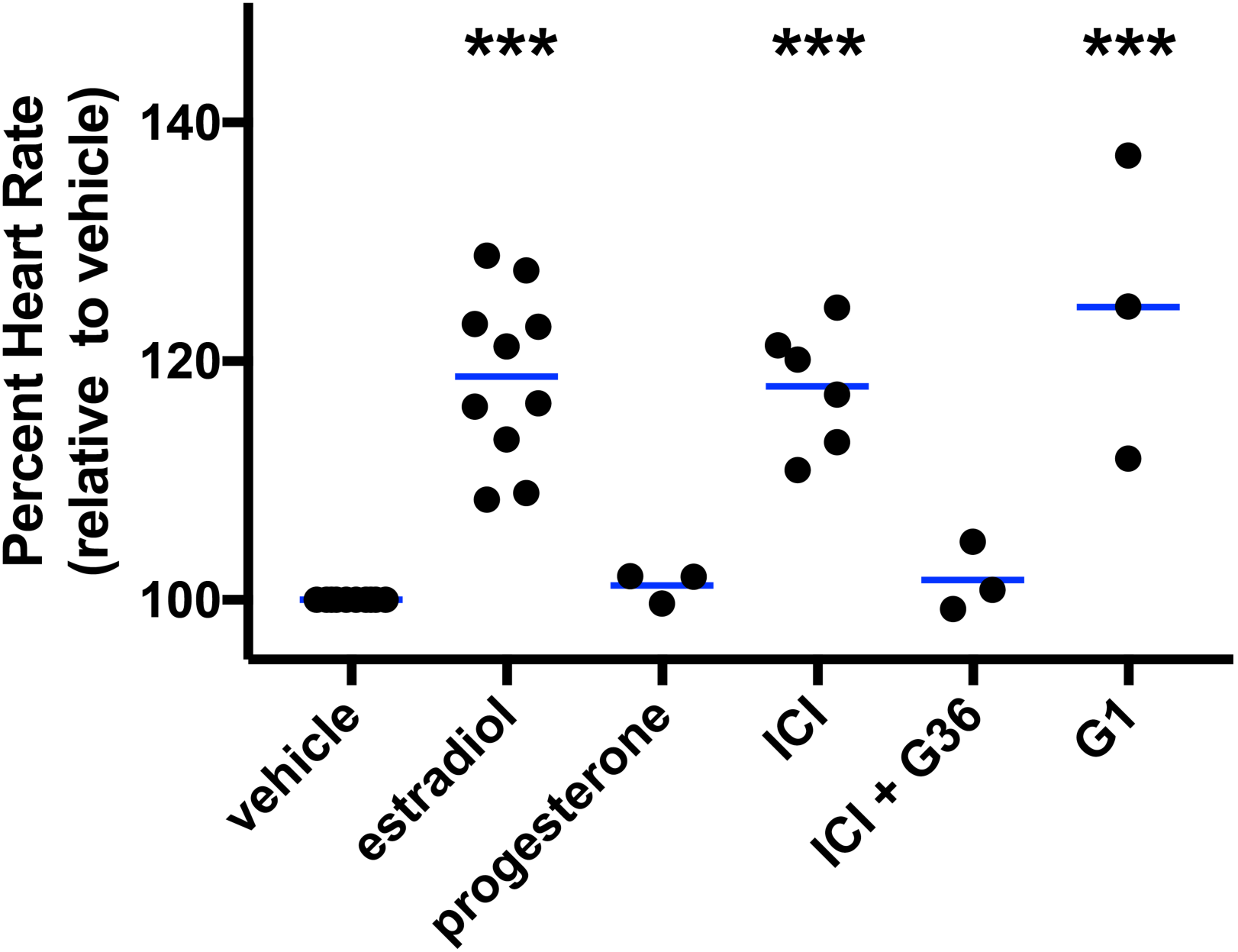
Estradiol and GPER agonists increased heart rate in zebrafish embryos. Wildtype embryos were incubated in water containing vehicle (0.1% DMSO), estradiol (3.67 μM, ER/GPER agonist), progesterone (1 μM), ICI (10 μM ICI182.780, ER antagonist/GPER agonist), G1 (1 μ M, GPER agonist), G36 (1 μM, GPER antagonist)or two chemicals in combination at 49 hours post fertilization and heart rates were measured 1 hour following treatment. ***, p<0.0001 compared to vehicle, ANOVA with Dunnett’s test. Each black circle represents the mean heart rate from a single clutch of embryos (3-16 embryos per clutch). Clutches in the same treatment group were assayed on different days. Horizontal blue lines are the mean of each treatment.

To explore whether heart rate was influenced by nuclear estrogen receptor or GPER signaling pathways, we employed a pharmacological approach. We exposed embryos to ICI182,780 (fulvestrant), a well-characterized ERα and ERβ antagonist (23) that also acts as a GPER agonist (8). Following one hour exposure to ICI182,780, heart rate was significantly increased (Fig. 1, mean difference in heart rate 29.81 ± 4.75 bpm). This effect was blocked by co-administration of G36, a specific GPER antagonist (24) (Fig. 1, mean difference in heart rate 4.10 ± 6.23 bpm), suggesting that estradiol increases heart rate via GPER. We also exposed embryos to G1, a specific GPER agonist with no detectable agonist activity against nuclear estrogen receptors (25), and found that heart rate increased significantly (Fig. 1, mean difference in heart rate 40.98 ± 6.35 bpm). Together, our pharmacological results suggest that GPER regulates heart rate acutely.

To definitively test the hypothesis that estradiol regulates heart rate via GPER, we generated GPER mutant embryos, exposed them to estrogen receptor modulators and assayed heart rate. Using CRISPR-Cas technology (26), we generated embryos with a 131 basepair deletion in the *gper* open reading frame (Fig. 2A, B). Embryos were viable and grossly normal, allowing us to measure heart rate (Fig. 2C, D). We exposed homozygous maternal zygotic *gper* mutant embryos (*MZgper-/-*) to estradiol or to ICI182,780 and found no increase in heart rate compared to embryos exposed to vehicle (Fig. 2E). Our results demonstrate that estradiol increases heart rate in a GPER-dependent manner. Note that zygotic *gper* mutants exhibited increased heart rate in response to estradiol (Fig. S1, mean difference in heart rate 29.11 ± 3.56 bpm), indicating that GPER is maternally deposited into oocytes and expressed in embryos. This is consistent with previously published results that detected *gper* transcript in zebrafish embryos at 1 hpf, suggesting the presence of maternally loaded *gper* mRNA (27).

**Figure 2.**
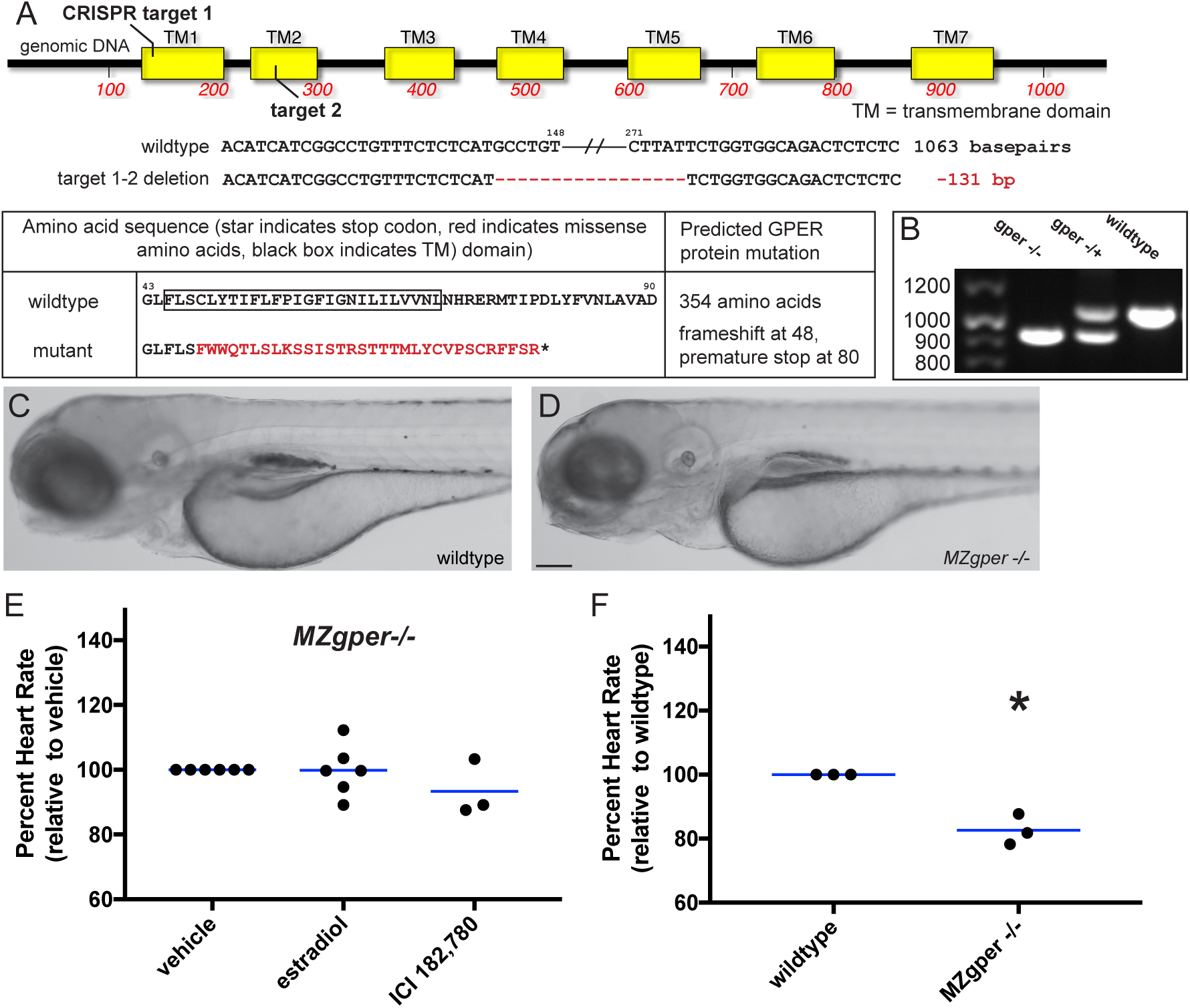
Abnormal heart rate in *gper* mutant zebrafish. **(A)** Genomic DNA of *gper*^*uab102*^ zebrafish contains a 131 basepair deletion in the *gper* coding region between CRISPR guide RNA targets 1 and 2, resulting in a premature stop codon in the GPER protein. Red dashes indicate DNA deletions, mutated amino acids are shown in red. **(B)** Genomic DNA was harvested from individual embryos, *gper* was PCR amplified and separated on an agarose gel to identify deletion mutations. **(C-D)** 3 day post fertilization wildtype and maternal zygotic *gper*^*uab102*^ homozygous larvae (*MZgper-/-*) exhibit similar gross morphology. Images are lateral views, anterior to the left, dorsal to the top. Scale bar, 500 μm. **(E)** Neither estradiol (ER/GPER agonist, 3.67 μM) nor ICI182,780 (ER antagonist/GPER agonist, 10 μM) changed heart rate significantly compared to vehicle (0.1% DMSO) in *MZgper-/-*, two-way ANOVA followed by F test, p=0.27. **(F)** *MZgper-/-* exhibited lower basal heart rate than age-matched wildtype embryos. *, p<0.05 compared to wildtype, paired t test. Each black circle represents the mean heart rate from a single clutch of embryos (≥ 7 embryos per clutch). Clutches in the same treatment group or genotype were assayed on different days. Horizontal blue lines are the mean of each treatment.

To test whether endogenous estrogens regulate heart rate during embryonic development, we examined basal heart rate in GPER mutant embryos reared in untreated water, reasoning that if heart rate was reduced, then that would suggest that endogenous estradiol regulates heart rate via GPER. We compared heart rate in wildtype versus *MZgper-/-* embryos at 50 hpf and found that *MZgper-/-* embryos had reduced heart rate compared to wildtype (Fig. 2F, mean difference in heart rate between wildtype and mutant -30.80 ± 7.07 bpm). These results demonstrate that GPER is required for normal basal heart rate in embryos and strongly suggest that endogenous estrogens influence heart rate via GPER.

Whether GPER acts as an autonomous estrogen receptor *in vivo* is controversial. Previous reports suggest that GPER activity might require interaction with nuclear estrogen receptors at the membrane or that estrogens activate GPER indirectly, by binding to nuclear receptors in the cytosol that then activate downstream proteins, including GPER (15, 28). To determine whether nuclear estrogen receptors influence heart rate, we generated zebrafish with loss-of-function mutations in each nuclear estrogen receptor gene: *esr1* (ERα), *esr2a* (ERβ1) and *esr2b* (ERβ2) (Fig. S2–S4). All mutant embryos were viable and grossly normal, allowing us to measure heart rate (Fig. S2–S4). To test whether estradiol increases heart rate via nuclear estrogen receptors, we exposed 49 hpf *esr1*^*-/-*^, *esr2a*^*-/-*^ and *esr2b*^*-/-*^ embryos to estradiol or vehicle for one hour and assayed heart rate. Following estradiol exposure, heart rate was increased in all mutants compared to vehicle control (Fig. 3A, mean difference in heart rate between estradiol and vehicle 25.04 ± 5.68 bpm for *esr1*^*-/-*^, 37.23 ± 7.66 bpm for *esr2a*^*-/-*^, 32.48 ± 1.92 bpm for *esr2b*^*-/-*^), similar to what we observed when wildtype embryos were exposed to estradiol (Fig. 1). These results demonstrate that nuclear estrogen receptors are not necessary for estradiol-dependent increase in heart rate.

**Figure 3.**
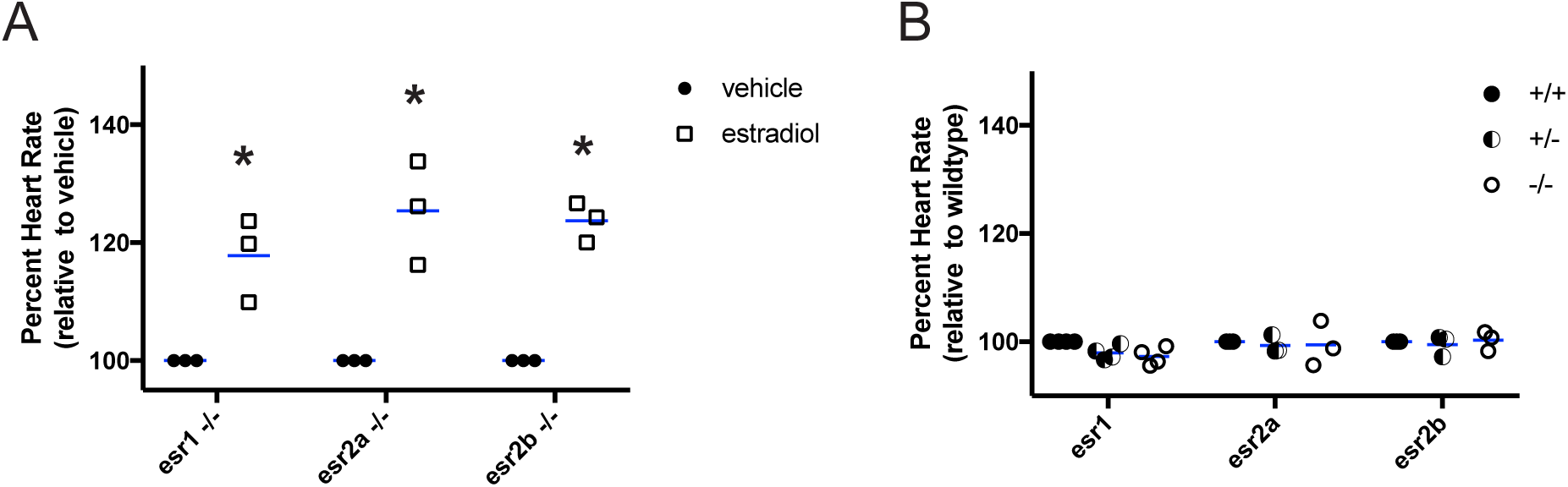
Normal heart rate in nuclear estrogen receptor mutants. **(A)** Homozygous mutant embryos at 49 hour post fertilization were incubated in water containing estradiol (ER/GPER agonist, 3.67 μM) or vehicle (0.1% DMSO) and heart rate was measured 1 hour post treatment. Estradiol increased heart rate compared to vehicle in zebrafish with homozygous mutations in ERα (*esr1 -/-*), ERβ1 (*esr2a -/-*), ERβ2 (*esr2b -/-*). *p<0.05 compared to vehicle within genotype, paired t-test. **(B)** Basal heart rate was measured at 50 hours post fertilization in embryos reared in untreated water. Heart rate was not significantly different in homozygous mutant (-/-) embryos compared to heterozygous (-/+) and wildtype (+/+) siblings for each *esr* mutant, two-way ANOVA. Each black circle represents the mean heart rate from a single clutch of embryos (4-8 embryos per clutch). Clutches in the same treatment group or genotype were assayed on different days. Horizontal blue lines are the mean of each treatment.

To test whether endogenous estrogens regulate heart rate via nuclear estrogen receptors, we bred heterozygous fish to generate embryos homozygous for mutations in either *esr1*, *esr2a* or *esr2b* genes and assayed heart rate in 50 hpf embryos. We observed no significant difference in basal heart rate between homozygotes, heterozygotes or wild type siblings within the same clutch (Fig. 3B, mean difference in heart rate between homozygote and wildtype -4.34 ± 1.37 for *esr1*, -0.46 ± 3.75 for *esr2a*, -2.37 ± 3.26 for *esr2b*; between heterozygote and wildtype -3.34 ± 1.02 for *esr1*, -0.91 ± 1.53 for *esr2a*, 0.63 ± 1.66 for *esr2b*). These results demonstrate that nuclear estrogen receptors are not required for the establishment of normal basal heart rate in embryos.

It is possible that the mutations generated in each nuclear estrogen receptor gene do not cause loss of functional estrogen receptor proteins. To exclude this possibility and show that *esr* mutants exhibit loss of functional ER proteins, we generated *esr* mutants on the *Tg(5xERE:GFP)*^*c262/c262*^ transgenic background, where green fluorescent protein (GFP) expression occurs in cells with activated nuclear estrogen receptors (5) (referred to as 5xERE:GFP). Previous studies using whole mount *in situ* hybridization demonstrated that *esr1* is expressed in embryonic heart valves while *esr2b* is expressed in the liver (6), therefore we hypothesized that mutants would fail to upregulate GFP in tissues where the relevant receptor is normally expressed. We exposed 2-3 day post fertilization (dpf) *5xERE:GFP*, *5xERE:GFP;esr1*^*-/-*^, *5xERE:GFP;esr2a*^*-/-*^ and *5xERE:GFP;esr2b*^*-/-*^ embryos to 100 ng/ml estradiol overnight and assayed fluorescence. Consistent with *esr* gene expression patterns, *5xERE:GFP;esr1*^*-/-*^ larvae exhibited fluorescence in the liver but not in the heart (Fig. S2), whereas *5xERE:GFP;esr2b*^*-/-*^ larvae exhibited fluorescence in the heart but not in the liver (Fig. S4). *esr2a* transcript was not detected at these embryonic and larval stages and, as expected, we saw no change in fluorescence between *5xERE:GFP* and *5xERE:GFP;esr2a*^*-/-*^ (Fig. S3). We conclude that the zebrafish nuclear estrogen receptor mutants lack estrogen receptor function.

Deleterious mutations can induce genetic compensation (29), however results from the 5xERE:GFP *esr* mutants suggest that compensatory expression of *esr* genes is not occurring. For example, it is possible that in the *esr1* mutant there is compensatory upregulation of *esr2a* and/or *esr2b* that masks a heart rate phenotype. If *esr2a* or *esr2b* were upregulated in *esr1* mutants, then we would expect to see fluorescence in the heart in *5xERE:GFP;esr1*^*-/-*^ embryos. Instead, we observed no fluorescence in the hearts of *5xERE:GFP;esr1*^*-/-*^ embryos (Fig. S2). Similarly, we observed no ectopic fluorescence in *5xERE:GFP;esr2b*^*-/-*^ embryos (Fig. S4), suggesting that *esr* genes are not compensating for one another in the multiple zebrafish *esr* mutants.

To further test whether nuclear estrogen receptor signaling is influenced by GPER, we generated *gper* mutants on the *5xERE:GFP* transgenic background and asked whether estradiol exposure reduced nuclear estrogen receptor activity in mutants compared to wildtype. Following overnight exposure to estradiol, 3 dpf *5xERE:GFP* and *5xERE:GFP;MZgper-/-* larvae exhibited similar fluorescence (Fig. S5). These results demonstrate that nuclear estrogen receptor transcriptional activity does not require GPER and support the hypothesis that GPER acts as an autonomous estrogen receptor *in vivo*.

Heart rate can be modulated by cardiomyocytes in the heart, or by cells in the central nervous system, which directly innervates the heart to modulate heart rate and also regulates the release of humoral factors, such as thyroid hormone, that bind to receptors in cardiomyocytes and regulate heart rate (30). To determine whether GPER regulates heart rate tissue autonomously, we performed whole mount *in situ* hybridization to test whether *gper* transcripts are expressed in 50 hpf zebrafish embryo hearts. We did not detect transcript in the heart or in the vasculature. In contrast, we detected *gper* mRNA in three discrete anatomic areas of the brain: the preoptic and olfactory areas and in the ventral hypothalamus-pituitary (Fig. 4A–C). Thus, *gper* localization is consistent with the hypothesis that GPER acts in the brain, and not through cells in the heart, to regulate heart rate.

**Figure 4.**
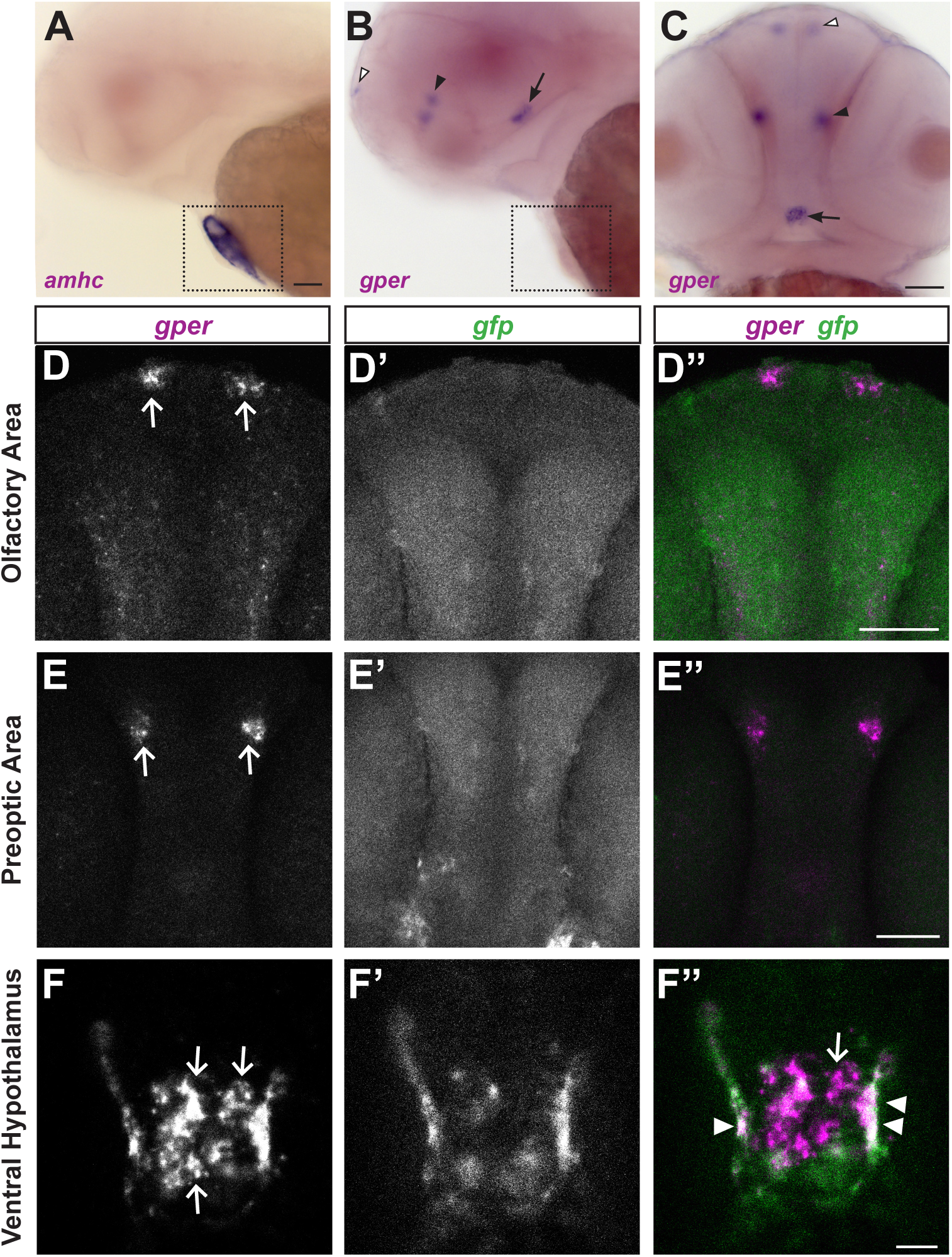
*gper* expression in the brain. **(A-C)** Whole mount colorimetric *in situ* hybridization was performed on wildtype embryos at 50 hours post fertilization (hpf). **(A)** *amhc* (alpha-myosin heavy chain) antisense RNA labels atrial myocardial cells in the heart (boxed). **(B, C)** *gper* antisense RNA labels a bilaterally symmetric cluster of cells in the olfactory area (white arrowheads) and preoptic area (black arrowhead) and a medial cluster of cells in the ventral hypothalamus (arrows). No label was detected in the heart. Lateral views with anterior to the left (A,B), ventral view with anterior to the top (C), scale bars = 100 μm. **(D-F)** Double fluorescent in situ hybridization performed on 48 hpf *Tg(5xERE:GFP)c262* embryos following overnight exposure to 100 ng/ml estradiol. *gfp* marks cells with active nuclear estrogen receptors. Confocal images of selected Z-slices (0.975 μm) show that *gper* is expressed in the olfactory area (D) and preoptic area (E) in cells lacking *gfp* (D”, E”, scale bars = 50 μm). In the ventral hypothalamus (F), *gper* is expressed in a medial cluster of cells lacking *gfp* (arrows, F, F”), whereas *gper* is expressed together with *gfp* more laterally (arrowheads, F”, scale bar = 10 μm). In merged images, *gper* is magenta, *gfp* is green and areas of colocalization are white. Dorsal views, anterior to the top.

Genetic evidence using *esr* mutants suggests that GPER acts independently of nuclear estrogen receptors to regulate heart rate (Fig. 3). To further test the hypothesis that GPER acts as an autonomous estrogen receptor *in vivo*, we asked whether GPER and nuclear estrogen receptors are expressed in the same cells in the brain, reasoning that if GPER and nuclear estrogen receptors fail to colocalize, this would support the idea that GPER acts as an autonomous estrogen receptor *in vivo*. We exposed 1 dpf 5xERE:GFP embryos to 100 ng/ml estradiol overnight. At 48 hpf, we fixed the embryos and used two color fluorescent in situ hybridization to detect *gfp* and *gper* transcripts simultaneously. Since all three nuclear estrogen receptor genes activate the 5xERE:GFP transgene, detecting *gfp* allows us to monitor activity of all three estrogen receptors using a single RNA probe. In the olfactory and preoptic areas, we found no colocalization between *gfp* and *gper* (Fig. 4D, E). In the ventral hypothalamus, we found a cluster of cells at the midline expressing *gper* but not *gfp*. Surrounding this region of *gper*-positive cells was a bilaterally symmetric ‘U’- shaped labeling pattern of cells expressing both *gper* and *gfp* (Fig. 4F). Thus, GPER and nuclear estrogen receptors are expressed in unique and overlapping cells in the brain, supporting the hypothesis that GPER can act independently of nuclear estrogen receptors *in vivo*.

At 2 dpf, the ventral hypothalamus and pituitary are contiguous, suggesting that gper could be expressed in both locations. The pituitary can regulate heart rate by secreting thyroid stimulating hormone (TSH), which stimulates the thyroid to produce thyroid hormone, which has been previously shown to increase heart rate in mammals (31–41). Thus, we hypothesized that GPER is expressed in pituitary cells that express thyroid stimulating hormone, where it regulates production of thyroid hormone. To test whether GPER is expressed in thyrotropic cells in the pituitary, we performed whole-mount two color fluorescent in situ hybridization to detect *gper* and thyroid stimulating hormone (*tshb*), a marker of thyrotropic cells (42). We identified a subpopulation of *tshb-*positive cells that also expressed *gper* (Fig. 5A), suggesting that GPER is expressed in thyrotropic pituitary cells.

**Figure 5.**
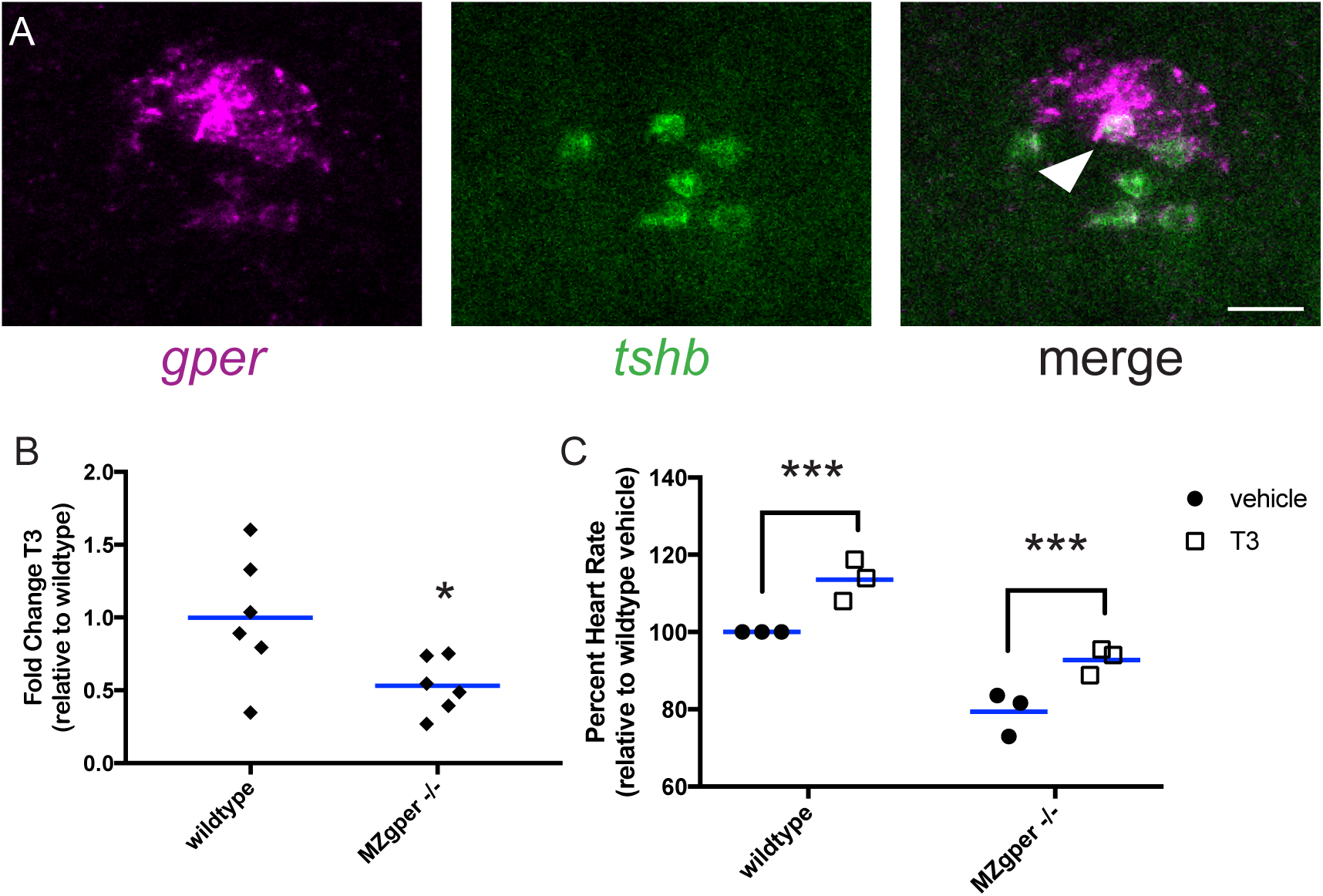
Triiodothyronine (T3) is reduced in *gper* mutant and rescues *gper* mutant heart rate phenotype. **(A)** Whole mount fluorescent in situ hybridization was used to detect *gper* and *thyroid stimulating hormone* (tshb) mRNA expression in wildtype 48 hpf embryos. White arrowhead indicates region of co-localization. Confocal images of single z-slices with anterior towards the top. Scale bar, 25 μm. (B) Triiodothyronine levels are reduced in maternal zygotic *gper*^*uab102*^ mutants compared to wildtype. Each black diamond represents mean results of sample duplicates from 50 pooled embryos at 49 hours post fertilization (hpf). * p<0.05, two-tailed t test. (C) T3 (5 nM) increases heart rate in wildtype and *MZgper -/-* mutant embryos compared to vehicle (0.1% methanol). ***p<0.0005 compared to vehicle within genotype, two-way ANOVA followed by F test. Each black circle represents the mean heart rate from a single clutch of embryos (≥ 6 embryos per clutch). Clutches in the same treatment group were assayed on different days. Horizontal blue lines are the mean of each treatment.

Since GPER is expressed in pituitary cells (Fig. 5A) and since the active form of thyroid hormone, triiodothyronine (T3), is known to be a humoral factor capable of regulating heart rate in mammals (31–41), we hypothesized that activation of GPER stimulates pituitary cells to increase levels of T3, thus leading to an increase in heart rate in zebrafish embryos. To test this hypothesis, we measured total T3 levels in *gper* mutant and wildtype embryos using an enzyme-linked immunosorbent assay (ELISA). At 49 hpf, *gper* mutant embryos showed a 50% reduction in total T3 levels (Fig 5B). While T3 is known to increase heart rate in mammals, whether this function is conserved in zebrafish embryos is not known. In adult zebrafish, a recent study reported that T3 may alter heart rate in a temperature specific manner, partially rescuing a decrease in heart rate after cold exposure (43). To determine whether T3 increases heart rate in zebrafish embryos, we exposed wildtype embryos to 5 nM T3 and found a mean 13% increase in heart rate compared to embryos exposed to vehicle (Fig. 5C; mean difference in heart rate 22.86 ± 3.43 bpm). We then tested whether T3 could rescue the reduced heart rate phenotype in *gper* mutant embryos. Following one hour exposure to 5 nM T3, *MZgper-/-* heart rate was increased 16%, to near wildtype levels (mean difference in heart rate 23.60 ± 4.98 bpm). This suggests that GPER is regulating T3 levels to maintain normal heart rate in zebrafish embryos.

## DISCUSSION

Here we provide evidence that estrogens signal through a non-canonical estrogen receptor, the G protein-coupled estrogen receptor (GPER), to regulate heart rate in zebrafish embryos by altering levels of thyroid hormone T3. Our results also support the hypothesis that GPER acts as an autonomous estrogen receptor *in vivo*. Previous reports using cultured cells demonstrated that fluorescently labeled or isotopic estradiol specifically binds membranes from cells expressing GPER (8, 9). Additionally, estradiol exposure increased cyclic AMP and calcium levels in HEK293 and COS7 cells in a GPER-dependent manner (8, 9), while estradiol exposure increased phosphoinositide 3-kinase activity in SKBR3 breast cancer cell line in a GPER-dependent manner (8). However, because these studies utilized cells that either express artificially high levels of GPER or are tumorigenic, the findings do not address whether GPER acts as an estrogen receptor *in vivo* under normal physiologic conditions. Our genetic and pharmacologic results strongly suggest that GPER is an estrogen receptor *in vivo*. If estradiol was binding to ERα or ERβ, and these receptors activated GPER, then we would expect to see no increase in heart rate in *esr1*, *esr2a* or *esr2b* mutants following exposure to estradiol. Instead, all *esr* mutants responded normally to estradiol exposure (Fig. 3), suggesting that ER and GPER signaling pathways are distinct in this context. Consistent with these results, we found *gper* transcript expressed in cells in the brain that lack nuclear estrogen receptor activity (Fig. 4), further supporting the hypothesis that GPER responds to estrogens independently of nuclear estrogen receptors *in vivo*. Studying the influence of estrogens on heart rate in zebrafish embryos is a powerful *in vivo* system where GPER activity is dissociated from classical nuclear estrogen receptor signaling.

Between 2 and 5 dpf, zebrafish heart rate normally increases (44, 45). Our results support the hypothesis that endogenous estradiol regulates this increase in heart rate. The finding that GPER mutant embryos have lower basal heart rate compared to wildtype embryos implicates endogenous estradiol. Additionally, a recent HPLC analysis of endogenous estradiol concentration in zebrafish embryos found that estradiol concentrations increased from 137 pg/embryo at 48 hpf to 170 pg/embryo at 72 hpf (46). Taken together, these results support the hypothesis that endogenous estradiol regulates heart rate in zebrafish embryos and larvae. The source of embryonic estradiol, whether synthesized by the embryo or maternally deposited in the yolk, is not known.

There are several mechanisms by which GPER activity in the brain could regulate heart rate, for example by modulating sympathetic and parasympathetic nerve activity or by regulating the release of humoral factors, such as thyroid hormone. Expression of *gper* transcript in thyrotropic cells in the pituitary (Fig 5A) and decreased levels of total T3 in *gper* mutants (Fig 5C) support the latter hypothesis. There are three primary mechanisms by which GPER could promote the increase of total T3: 1) by increasing levels of thyroid stimulating hormone, leading to release of thyroxine (T4) from the thyroid, which is then converted to T3, 2) by increasing the conversion of T4 to T3, or 3) by blocking the conversion of T3 into inactive metabolites, such as 3,5-Diiodo-L-thyronine (T2) and reverse T3 (RT3; 3,3’5’-triiodothyronine). *gper* expression in *tshb*-positive cells in the pituitary supports the first hypothesis. In humans, thyroid stimulating hormone is thought to be required for the differentiation of the thyroid (47). Curiously, zebrafish mutant embryos that lack thyrotrope progenitor cells and *tsh* gene expression still produce thyroid follicle cells and T4 (48). This suggests that thyrotropes may not be required to produce T3, however total T3 was not measured in this study. It is possible that the localized production of T4 is sufficient to stimulate thyroid development, while total T3 is reduced. In support of this, a majority of the cartilage in the pharynx is missing in thyrotrope deficient zebrafish (48), which suggests that even though the thyroid begins to differentiate, tissues adjacent to the thyroid do not, presumably due to reduced total T4 in circulation. While TSH signaling may not be required for the development of the thyroid, it may be required for secondary functions, including proper regulation of heart rate.

We cannot exclude the possibility that GPER activity leads to increased expression or activity of deiodinase enzymes that convert T4 to T3, or that GPER activity reduces the expression or activity of enzymes that metabolize T3. All four deiodinases genes (*dio1*, *dio2*, *dio3a* and *dio3b*) are expressed in zebrafish as early as 24 hpf (49) and are therefore available to convert thyroid hormones at 48 hpf, when we observe changes in heart rate. Interestingly, at 24 hpf *dio2*, the enzyme that converts T4 to T3, is expressed in the pituitary (50) in addition to its expression in the thyroid. *dio3a*, which inactivates T3 by conversion to RT3 and T2, is also expressed in the brain and thyroid at 24 hpf (50). Previous work suggests that GPER influences neurotransmitter release and cAMP levels (51). cAMP was shown to increase deiodinase activity in the brain leading to increased T3 levels (52–54). Therefore, GPER activity could trigger neuronal activity that leads to increased activity of deiodinases and increased production of T3, independently of TSH, to regulate heart rate (30).

More generally, it is not known to what extent estrogens influence thyroid hormone signaling. Our work suggests that endogenous estrogens influence T3 levels. Previous work suggests that environmental estrogens may also influence thyroid hormone signaling. Exposure to the plasticizer diethylhexyl phthalate (DEHP) increased total T3 levels in zebrafish larvae and upregulated thyroid signaling genes thyroglobulin (*tg*), transthyretin (*ttr*), and *dio2* (55). DEHP exhibits estrogen-like activity, although the receptor by which it acts has not been determined. DEHP can inhibit tamoxifen induced apoptosis and also induce cell proliferation in GPER positive MCF-7 cells, but not in GPER negative MDA-MB-231 cells (56), suggesting that DEHP can activate GPER. One possibility is that DEHP increases T3 levels in zebrafish larvae via GPER activation. Similarly, chronic exposure to perfluorooctanesulphonic acid (PFOS), a surfactant that enhances the effects of estradiol (57), increased total T3 levels in juvenile zebrafish and upregulated thyroid signaling genes thyroid hormone receptor β, the sodium/iodide symporter *slc5a5*, *dio1* and *dio2* (58). We speculate that like estradiol, the environmental endocrine disruptors DEHP and PFOS modulate T3 levels by activating GPER. This raises the important consideration that diverse environmental estrogens could alter thyroid signaling and thus cardiac function.

While our results illuminate GPER signaling in the context of embryonic heart rate, it is not clear to what extent GPER influences heart rate at later stages of development. At larval, juvenile and adult stages it is impossible to assess heart rate without immobilizing or anesthetizing zebrafish, manipulations that themselves influence heart rate. In adult mice with mutations in GPER, there was no significant difference in basal heart rate between mutant and wildtype mice of either sex (16, 17, 59). It is possible that GPER regulates heart rate in embryos but not in adults. Additionally, heart rate in GPER mutant mice was assayed using general anesthesia, which is known to depress heart rate compared to conscious mice (60). Anesthesia may mask the effect of GPER on basal heart that we observe in conscious animals. We also cannot exclude the possibility that the effects of GPER on heart rate are specific for zebrafish.

Considering that GPER deficient zebrafish embryos have altered T3 levels, it will be interesting to examine whether this deficiency exists at later developmental stages and whether GPER mutant adults have growth and metabolic defects consistent with reduced total T3. In *MZgper-/-* embryos, we observed no gross morphological detects up to 2 dpf, while mutant adults are viable and fertile. Zebrafish mutants with ∼70-90% reduced total T3 levels due to genetic ablation of *dio2* exhibit delayed swim bladder inflation, altered locomotor activity through 7 dpf, delayed fertility, reduced number of eggs, and reduction in viable fertilized eggs (61). In GPER deficient fish, the reduction in T3 levels is less drastic and we anticipate seeing less severe phenotypes as a result of the more modest decrease in T3.

In summary, this study identified a role for GPER in the regulation of embryonic heart rate via increased T3 levels. The zebrafish estrogen receptor mutants we developed enable experiments to rapidly and conclusively identify the causative estrogen receptor associated with any estrogen signaling phenotype, as demonstrated with the estradiol-dependent increase in heart rate reported here. This has significant implications for studies of estrogenic environmental endocrine disruptors, which are frequently tested on zebrafish to identify effects on embryonic development, organ formation and function (62). Zebrafish estrogen receptor mutants can now be used to determine whether such effects are specific for estrogen receptors and to identify the precise receptor target. Our results also establish a need to consider the impact on cardiac function when considering the toxicity of estrogenic environmental endocrine disruptors.

## Materials and Methods

### Zebrafish

Zebrafish were raised at 28.5°C on a 14-h light, 10-h dark cycle in the UAB Zebrafish Research Facility in an Aquaneering recirculating water system (Aquaneering, Inc., San Diego, CA). Wildtype zebrafish were AB strain (63) and all mutant and transgenic lines were generated on the AB strain. To visualize nuclear estrogen receptor activity, transgenic line *Tg(5xERE:GFP)*^*c262/c262*^ was used for all studies unless otherwise mentioned (5). All procedures were approved by the UAB Institutional Animal Care and Use Committee.

### Embryo collection

Embryos were collected during 10 minute intervals to ensure precise developmental timing within a group. Embryos were placed in Petri dishes containing E3B (60X E3B: 17.2g NaCl, 0.76g KCl, 2.9g CaCl_2_-2H_2_O, 2.39g MgSO_4_ dissolved in 1 liter Milli-Q water; diluted to 1X in 9 liter Milli-Q water plus 100 μL 0.02% methylene blue) and placed in an incubator at 28.5°C on a 14-h light, 10-h dark cycle. At 24 hours post fertilization (hpf), embryos were incubated in E3B containing 200 μM 1-phenyl 2-thiourea (PTU) to inhibit pigment production (63). Between 24 and 48 hpf, embryos were manually dechorionated and randomly divided into control and experimental treatment groups (10 to 30 embryos per treatment group) in 60mm Petri dishes and kept at 28.5°C until 49 hpf.

### Embryo treatments

At 49 hpf, embryos were incubated in E3B with estrogen receptor modulator(s) at 28.5°C for 1 hour. Estrogen receptor modulator treatments consisted of: 3.67 μM E2 (17β-estradiol; Sigma E8875, purity ≥ 98%), 10 μM ICI182,780 (fulvestrant; Sigma I4409, purity >98%), 1 μM G1 (Azano AZ0001301, purity ≥ 98%), 1 μM G36 (Azano, AZ-0001303, purity ≥ 98%), 1 μM progesterone (Sigma P0130, purity ≥ 99%), 5 nM 3,3’,5-Triiodo-L-thyronine (T3; Sigma T2877, purity ≥ 95%), vehicle (0.1% dimethylsulfoxide (DMSO), Fisher D128-500; purity ≥ 99.9% or 0.1% methanol, Fisher A411-4). All chemical stocks were made in 100% DMSO at 1000x and diluted in E3B embryo media to final concentration at the time of treatment, except for T3 which was made in 100% methanol for chemical stocks but was diluted in E3B media as above. The T3 concentration used was previously shown to be effective at inducing thyroid hormone-dependent gene expression in zebrafish (49). For rescue experiments (ICI182,780 + G36), final DMSO concentration was 0.2%. There was no difference in heart rate between embryos incubated in 0.1% or 0.2% DMSO (not shown). All vehicle controls shown in figures are 0.1% DMSO, except where indicated.

### Measurement of heart rates

All embryos were reared at 28.5 °C and heart rate was measured at room temperature. Following one hour incubation in treatment compounds at 28.5^o^C, heart rate (beats per minute, bpm) was calculated by counting the number of heart beats in fifteen seconds and multiplying that number by four. Prior to measurements, each dish was removed from the incubator and placed under the microscope light for 4 minutes at room temperature, allowing embryos to acclimate to the light and eliminate any effect of the startle response. Control groups were counted first and last to ensure that the overall heart rate did not increase during the duration of counting due to natural increases in heart rate during development. All heart rates were measured on a Zeiss Stemi 2000 dissecting microscope with a halogen transmitted light base (Carl Zeiss Microimaging, Thornwood, NJ).

### Generation of guide RNA and Cas9 mRNA

Plasmids pT7-gRNA and pT3TS-nCas9n were obtained from Addgene (numbers 46759, 46757) (26). pT7-gRNA was digested simultaneously with BsmBI, BglII and SalI for one hour at 37 °C followed by one hour at 55 ^°C^. To generate esr2a, esr2b and gper gRNAs, oligonucleotides containing target site sequences (see table below) were synthesized by Invitrogen. Oligos were hybridized to each other using NEBuffer3 restriction enzyme buffer (New England Biolabs) to generate double stranded target DNA and annealed into digested pT7-gRNA using Quick T4 DNA Ligase (New England Biolabs) as previously described (26). Guide RNAs were synthesized using the MegaShortScript T7 Kit (Life Technologies) using the relevant modified pT7-gRNA vector linearized with BamHI as a template. Guide RNA was purified using the RNA clean & concentrator kit (Zymo Research). To generate *esr1* guide RNA, target-specific oligonucleotides containing the SP6 (5’-ATTTAGGTGACACTATA) promoter sequence, a 20 base target site without the PAM, and a complementary region were annealed to a constant oligonucleotide encoding the reverse-complement of the tracrRNA tail as described (64). This oligo was used as a template for in vitro transcription using the MegaShortScript Sp6 Kit (LifeTechnologies). To generate *Cas9* mRNA, the pT3TS-nCas9n plasmid was linearized with XbaI and transcribed using the mMessage mMachine T3 kit (Life Technologies) and purified using RNA clean & concentrator kit (Zymo Research). RNA concentration was quantified using a Nanodrop spectrophotometer (Nanodrop ND-1000, ThermoFisher).

Target site sequences for *gper*, *esr1*, *esr2b* and *esr2a* oligonucleotides:

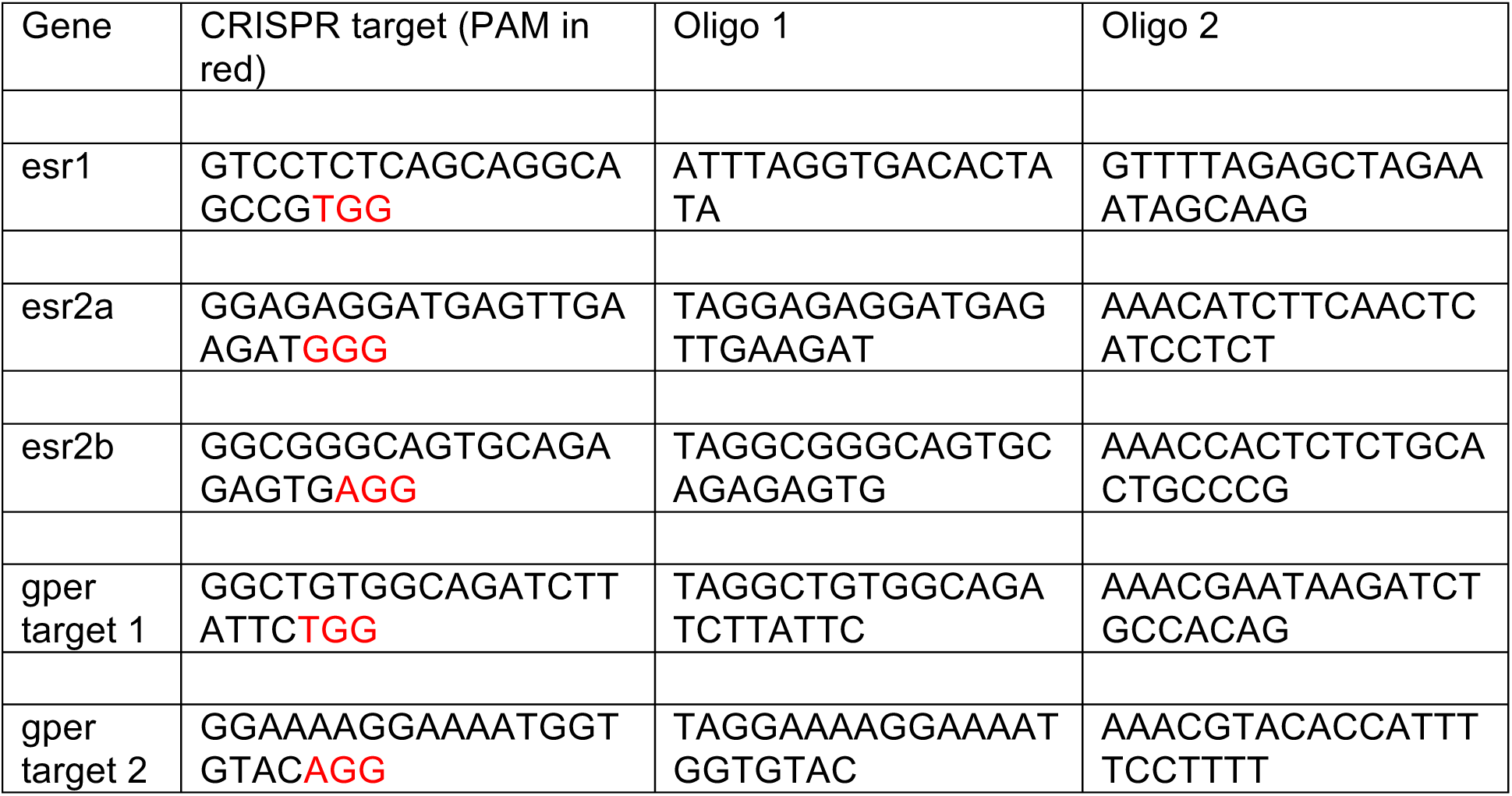

### Embryo injections

One-cell-stage embryos were injected using glass needles pulled on a Sutter Instruments Fleming/Brown Micropipette Puller, model P-97 and a regulated air-pressure micro-injector (Harvard Apparatus, NY, PL1–90). Each embryo was injected with a 1 nl solution of 150 ng/μl of Cas9 mRNA, 50 ng/μl of gRNA and 0.1% phenol red. Mixtures were injected into the yolk of each embryo. Injected embryos were raised to adulthood and crossed to wildtype fish (either AB or *Tg5xERE:GFP*^*c262*^) to generate F1 embryos. F1 offspring with heritable mutations were sequenced to identify loss of function mutations.

### Genomic DNA isolation

Individual embryos or tail biopsies from individual adults were placed in 100 μL ELB (10 mM Tris pH 8.3, 50 mM KCl, 0.3% Tween 20) with 1 μL proteinase K (800 U/ml, NEB) in 96 well plates, one sample per well. Samples were incubated at 55°C for 2 hours (embryos) or 8 hours (tail clips) to extract genomic DNA. To inactivate Proteinase K, plates were incubated at 98°C for 10 minutes and stored at -20°C.

### High resolution melt curve analysis

PCR and melting curve analysis was performed as described (65). PCR reactions contained 1 μl of LC Green Plus Melting Dye (BioFire Diagnostics), 1 μl of Ex Taq Buffer, 0.8 μl of dNTP Mixture (2.5 mM each), 1 μl of each primer (5 μM), 0.05 μl of Ex Taq (Takara Bio Inc), 1 μl of genomic DNA, and water up to 10 μl. PCR was performed in a Bio-Rad C1000 Touch thermal cycler, using black/white 96 well plates (Bio-Rad HSP9665). PCR reaction protocol was 98°C for 1 min, then 34 cycles of 98°C for 10 sec, 60°C for 20 sec, and 72°C for 20 sec, followed by 72°C for 1 min. After the final step, the plate was heated to 95°C for 20 sec and then rapidly cooled to 4°C. Melting curves were generated with either a LightScanner HR 96 (Idaho Technology) over a 70–95°C range and analyzed with LightScanner Instrument and Analysis Software (V. 2.0.0.1331, Idaho Technology, Inc, Salt Lake City, UT), or with a Bio-Rad CFX96 Real-Time System over a 70–95°C range and analyzed with Bio-Rad CFX Manager 3.1 software.

### Live imaging

Live zebrafish embryos and larvae were visualized using a Nikon MULTIZOOM AZ100 equipped with epi-fluorescence and an Andor Clara digital camera unless otherwise noted. To validate mutants with 5xERE reporter activity, larvae were treated overnight with 100 ng/mL estradiol beginning at 2-3 dpf. Following overnight treatment, larvae were washed in E3B, anesthetized with 0.04% tricaine and imaged in Petri dish containing E3B. For Fig. S1 H-K, larvae were mounted in bridged coverslips in E3B with 0.04% tricaine (63). Images were captured on a Zeiss Axio Observer.Z1 fluorescent microscope equipped with an Axio HRm camera and Zen Blue 2011 software (Carl Zeiss Microscopy, Oberkochen, Germany). Adjustments, cropping and layout were performed using Photoshop CS6 and InDesign CS6 (Adobe Systems Inc., San Jose, CA).

### RNA in situ hybridization

For synthesis of RNA probes, full-length *gper* open reading frame was amplified by PCR from genomic DNA extracted from 3 dpf larvae (*gper* coding region is within a single exon and therefore the open reading frame sequence is identical in genomic and cDNA) using primers 5’-ATGGAGGAGCAGACTACCAATGTG-3’ and 5’- CTACACCTCAGACTCACTCCTGACAG-3’.

For *tshb* probe, a 252bp product was amplified by PCR from cDNA (prepared from total RNA from 5 dpf AB larvae using RETROscript reverse transcription kit (ThermoFisher Scientific) with oligo(dT) primers) using primers 5’ GAGTTGGTGGGTCCTCGTTT 3’ and 5’ TGCTTGGGCGTAGTTGTTCT 3’. Each product was then TA cloned into pCR2.1 vector (Invitrogen). *amhc* and *gfp* probes were used as described (5, 66). All clones were verified by sequencing. Digoxigenin-labeled antisense RNA and FITC-labeled antisense RNA were transcribed using T7 and T3 polymerase, respectively, as previously described (5).

Colorimetric whole-mount *in situ* hybridization was performed on zebrafish embryos and larvae as described previously, using 5% dextran in the hybridization buffer (67, 68). Following colorimetric *in situ* hybridization, embryos were sequentially cleared in glycerol (25%, 50%, 75% in phosphate buffered saline), mounted in 4% low-melting temperature agarose, and imaged using a Zeiss Axio Observer.Z1 microscope with Zeiss Axio MRc5 camera and Zen Blue 2011 software. Fluorescent *in situ* hybridization (FISH) was performed as previously described (68) with the following modifications: After rehydration, Proteinase K treatment was extended to 35 minutes. Following hybridization, embryos were washed in 2xSSC prior to being placed in PBT. Embryos were blocked in 2% Roche blocking reagent in 100 mM Maleic acid, 150 mM NaCl, pH 7.5 (69). For double labeling, following development of anti-DIG-POD antibody, reaction was inactivated in 100 mM glycine pH 2 for 10 minutes then incubated in anti-FITC antibody. Following florescent *in situ* hybridization, embryos were cleared in 50% glycerol, mounted on a bridged coverslip and imaged using a Nikon A1/R scanning confocal microscope with Nikon Advanced Elements software.

### Measuring T3 levels

T3 levels were measured using enzyme-linked immunosorbent assay as previously described (70), with minor modifications, using T3 ELISA Kit (IBL America IB19107). Briefly, 50 embryos were pooled in 50 μl of PBS and pulsed sonicated intermittently for 5 mins, alternating 5 sec sonication and 5 seconds on ice, then vortexed intermittently for 10 mins, alternating 30 sec vortexing and 30 sec on ice. Samples where then centrifuged for 10 mins at 15,000g at 4°C. Supernatant was collected and diluted 1:8 in PBS. 50 μl was used per reaction in accordance with the manufacturer’s instructions. Each sample was tested in duplicate and the mean of duplicates were compared statistically.

### Experimental design and data analysis

Heart rate assays were conducted in separate experiments. Each experiment included comparing groups (treated vs untreated or mutant vs wildtype) using at least 3 embryos per group with all embryos from the same clutch. All experiments were replicated at least 3 times (n≥3) using different clutches on different days. This is essentially a complete block design with clutch/day as block. Mean heart rate of individual embryos from a clutch was used for comparing treatment groups (or mutant groups) within experiments using two-way ANOVA controlling for clutch/day effect. The overall treatment effect (or the genotype effect in some experiments) was tested using F test. If it was significant, Dunnett’s test was then used to compare each treatment group with the vehicle group or mutant group with the wildtype group. For some individual pairs of comparisons, paired t test was used. Significance level is 0.05. All the analyses were conducted using R (version 3.0.2). Graphs were produced using GraphPad Prism 7.0a software.

## Acknowledgements

We thank J.L. King and J.P. Souder for technical assistance and S.C. Farmer and the staff of the UAB zebrafish facility for animal care.

## Author contributions

S.N.R., H.E.E. and D.A.G. performed experiments and analyzed experimental data. X.C. performed statistical analyses of heart rate data. S.N.R. and D.A.G. conceived the project, D.A.G. supervised the project. All authors contributed to writing the paper and read and approved the final manuscript.

---

**Figure S1.**
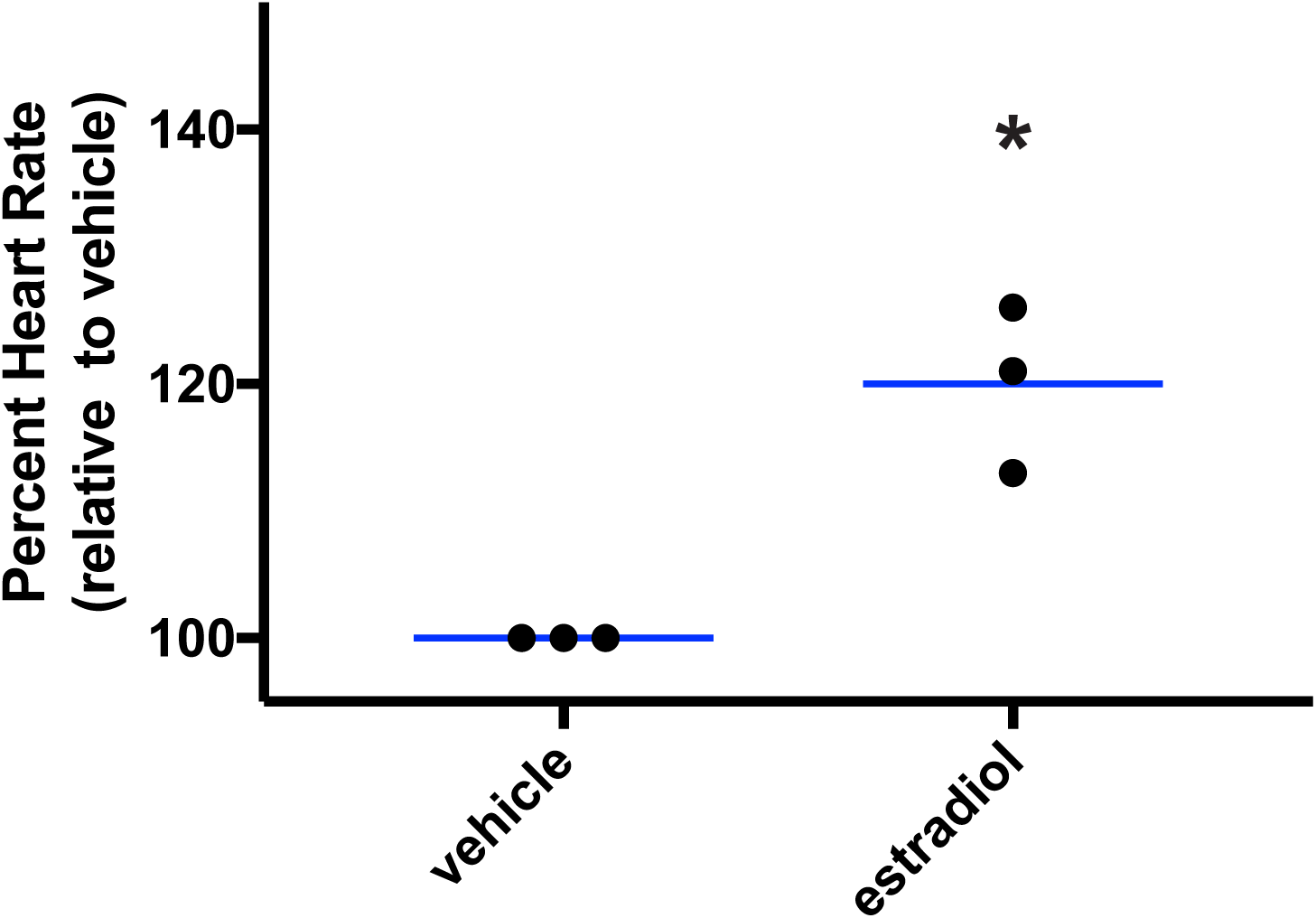
Zygotic *gper* mutant embryos are sensitive to estradiol. Zygotic homozygous *gper* mutant embryos were incubated in water containing estradiol (ER/GPER agonist, 3.67 μM) or vehicle control (0.1% DMSO) at 49 hours post fertilization and heart rates were measured 1 hour post treatment. *, p<0.05 compared to vehicle, paired t test. Each black circle represents the mean heart rate from a single clutch of embryos (≥ 6 embryos per clutch). Clutches in the same treatment group were assayed on different days. Horizontal blue lines are the mean of each treatment.

**Figure 2.**
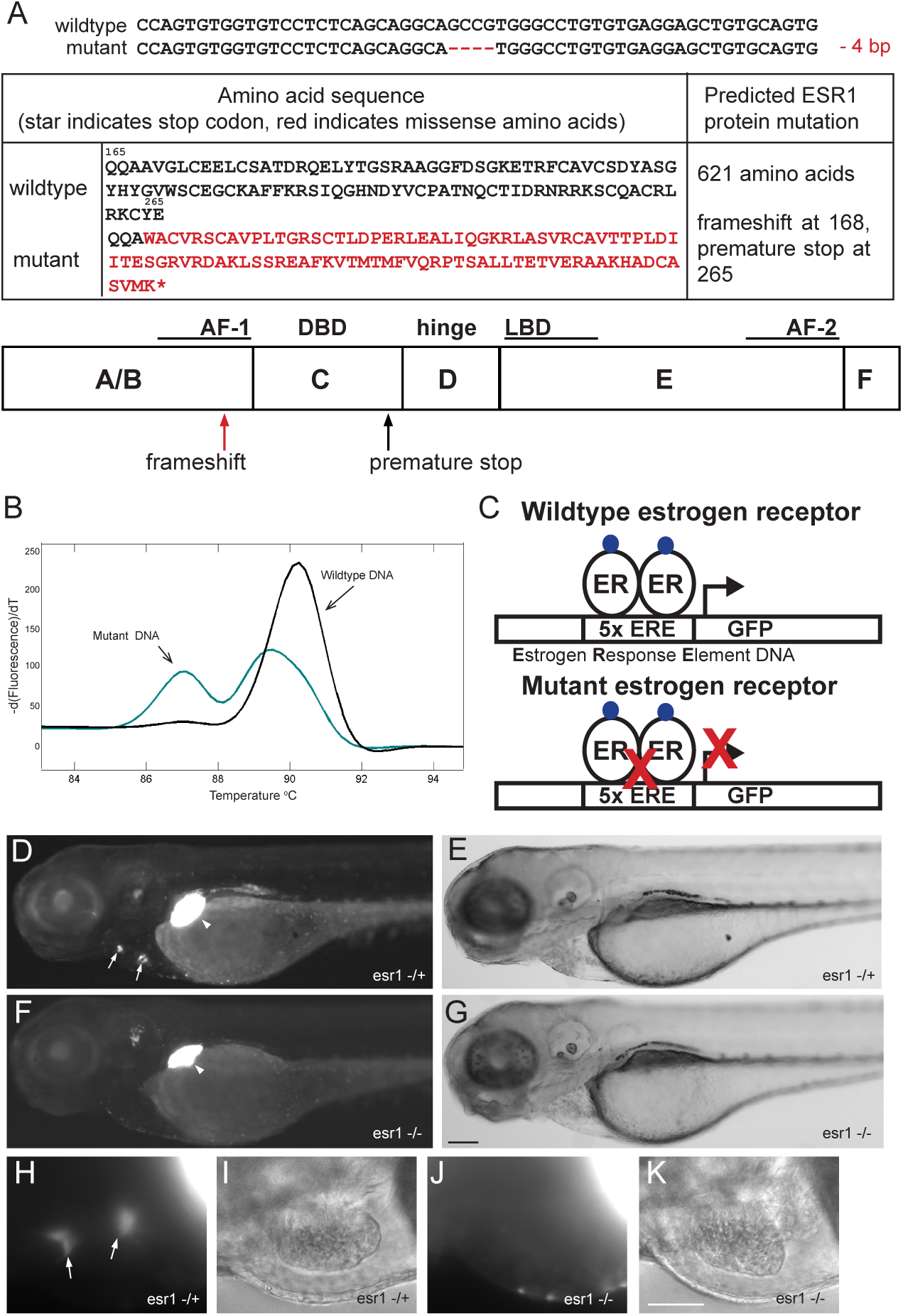
Generation and validation of *esr1* mutant zebrafish. **(A)** Genomic DNA of *esr1*^*uab118*^ zebrafish contains a 4 basepair deletion in the *esr1* coding region, resulting in a premature stop codon in the Esr1 (ERα) protein. Nucleotide deletions are shown as red dashes, amino acid mutations are in red. Map indicates site of frameshift mutation and premature stop codon (AF-1, activating function 1 domain; DBD, DNA binding domain; LBD, ligand binding domain; AF-2, activating function 2 domain). **(B)** High resolution melting curve analysis was used to distinguish mutants from wildtype. Curves represents DNA amplified from a wildtype AB (black) or *esr1*^*uab118*^ mutant zebrafish (cyan). **(C)** Strategy for validating zebrafish estrogen receptor mutants using transgenic 5xERE:GFP zebrafish. Mutants were generated on a transgenic background where estrogen receptor (ER) transcriptional activity is marked by green fluorescent protein (GFP) expression. Following exposure to estradiol, loss-of-function mutants should exhibit reduced fluorescence in cells expressing *esr1*. **(D-K)** 2-day post fertilization embryos were exposed to 367 nM (100 ng/mL) estradiol, live fluorescent images (D, F, H, J) and corresponding brightfield images (E, G, I, K) were taken at 3 d. *5xERE:GFP*^*c262;*^*esr1*^*uab118*^ homozygous larvae (*esr1 -/-*) exhibit normal morphology, but lack fluorescence in heart valves, wheres heterozygotes (*esr1 -/+*) exhibit fluorescent heart valves. High magnification images of the heart are shown in H-K. Arrows indicate heart valves, arrow head indicates liver. Images are lateral views, anterior to the left, dorsal to the top. Scale bars, 500 μm (D-G), 100 μm (H-K).

**Figure S3.**
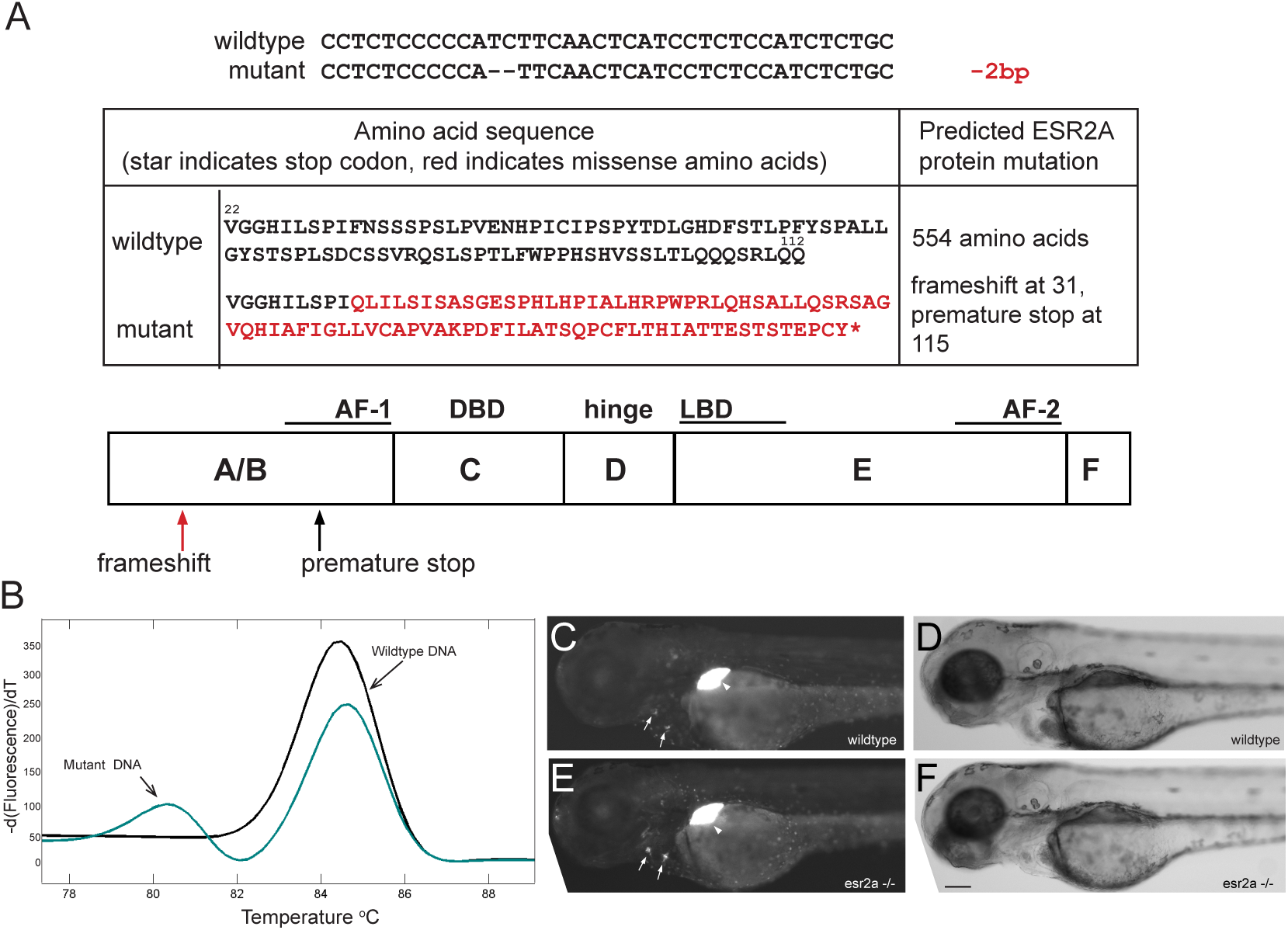
Generation of *esr2a* mutant zebrafish. **(A)** Genomic DNA of *esr2a*^*uab134*^ zebrafish contains an 2 basepair deletion (red dashes) in the *esr2a* coding region, resulting in a premature stop codon in the Esr2a (ERβ1) protein. Amino acid mutations are in red. Map indicates frameshift mutation and premature stop codon in the Esr2a protein. AF-1, activating function 1 domain; DBD, DNA binding domain; LBD, ligand binding domain; AF-2, activating function 2 domain. **(B)** High resolution melting curve analysis was used to distinguish mutants from wildtype. Curves represent DNA amplified from a wildtype AB (black) or *esr2a*^*uab134*^ mutant zebrafish (cyan). **(C-F)** *5xERE:GFP*^*c262*^ *and 5xERE:GFP*^*c262;*^*esr2a*^*uab134*^ (*esr2a -/-*) 3-day post fertilization (d) larvae were exposed to 367 nM (100 ng/mL) estradiol. Live fluorescent images (C, E) and corresponding brightfield images (D, F) were captured at 4 d. *esr2a -/-* larvae exhibit normal morphology and fluorescence, consistent with data demonstrating that *esr2a* is not expressed during these developmental stages. Arrows indicate heart valves, arrow head indicates liver. Images are lateral views, anterior to the left, dorsal to the top. Scale bar, 500 μm

**Figure S4.**
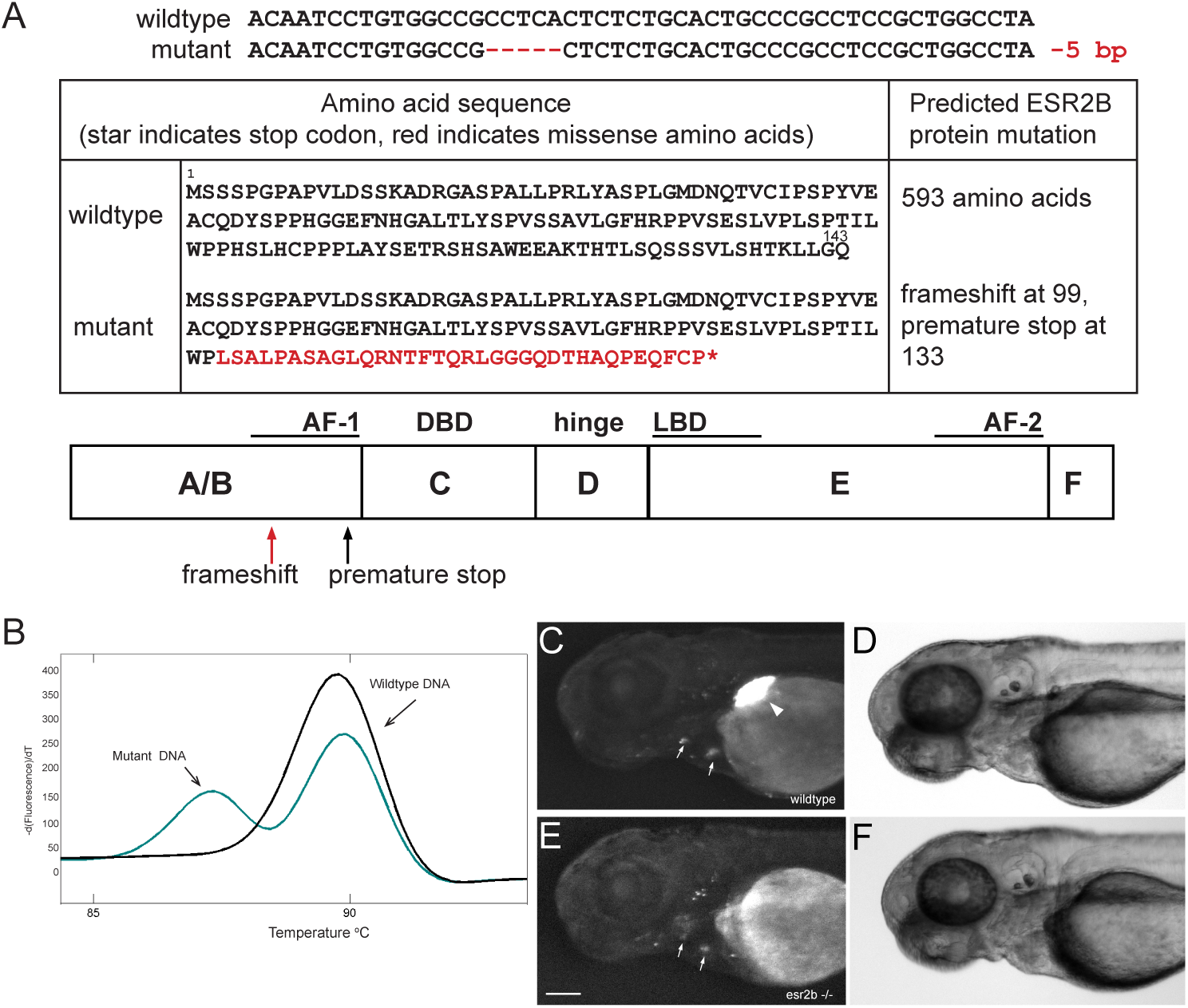
Generation and validation of *esr2b* mutant zebrafish. **(A)** Genomic DNA of *esr2b*^*uab127*^ zebrafish contains a 5 basepair deletion (red) in the *esr2b* coding region, resulting in a premature stop codon in the Esr2b (ERβ2) protein. Amino acid mutations are in red. Map indicates frameshift mutation and premature stop codon in the Esr2b protein. AF-1, activating function 1 domain; DBD, DNA binding domain; LBD, ligand binding domain; AF-2, activating function 2 domain. **(B)** High resolution melting curve analysis was used to distinguish mutants from wildtype. Curves represents DNA amplified from a wildtype AB (black) or *esr2b*^*uab127*^ mutant zebrafish (cyan). **(C-F)** *5xERE:GFP*^*c262;*^*esr2b*^*uab127*^ 3-day post fertilization (d) larvae were exposed to 367 nM (100 ng/mL) estradiol. Live fluorescent images (C, E) and corresponding brightfield images (D, F) were captured at 4 d. *5xERE:GFPc262;esr2buab127* homozygous larvae (*esr2b -/-*) exhibit normal morphology, but lack fluorescence in the liver. Arrows indicate heart valves, arrow head indicates liver. Images are lateral views, anterior to the left, dorsal to the top. Scale bar = 100 μm.

**Figure S5.**
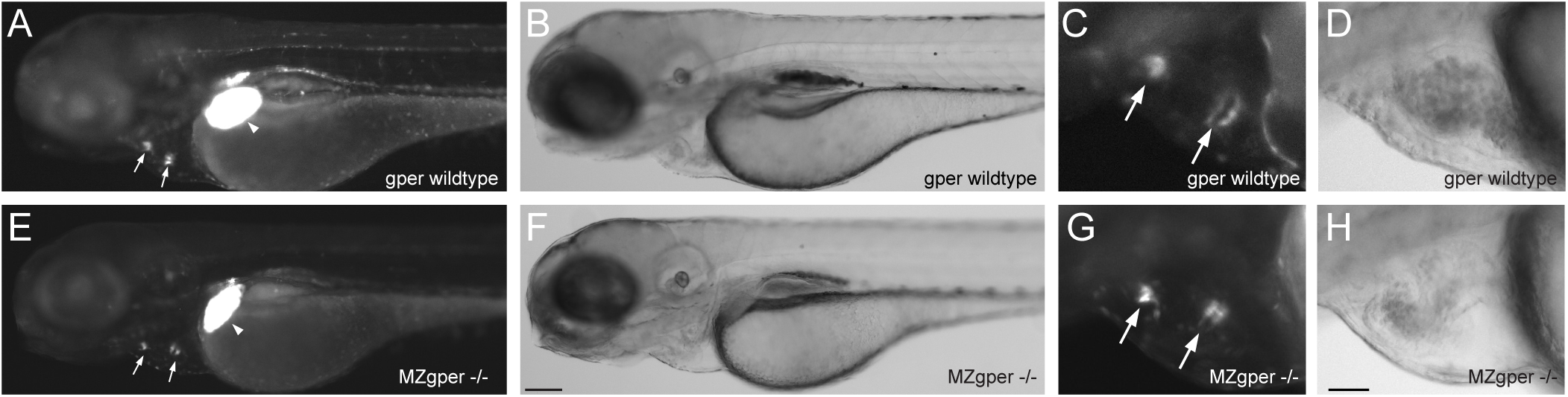
Nuclear estrogen receptor transcriptional activity is normal in *gper* mutant zebrafish. **(A-H)** Maternal zygotic *gper*^*uab102*^ homozygous larvae on the *5xERE:GFP*^*c262*^ transgenic background (*MZgper-/-*) were exposed to 367 nM (100 ng/mL) estradiol at 2-days post fertilization (2 d). Fluorescence (A, C, E, G) and corresponding brightfield images (B, D, F, H) were taken at 3 d. Fluorescence in the heart valves (arrows) and liver (arrow heads) is similar between MZ*gper -/-* and wildtype larvae. C, D, G, H, High magnification images of heart. Images are lateral views, anterior to the left, dorsal to the top. Scale bars, 500 μm (C-F), 100 μm (G-J).

